# Lead (Pb) Inhibits-PMA Induced Microglia Multinucleation

**DOI:** 10.1101/2023.04.04.535606

**Authors:** Stacia M. Nicholson, Karie Chen, Francis A.X. Schanne

## Abstract

Multinucleated Microglia are formed in response to aging, inflammation, the presence of some pathogens, and are a histological feature of several CNS pathologies. Multinucleated microglia may be involved with increased clearance of CNS debris and whole cells. The present study has sought to determine the best model by which to study multinucleation in microglia *in vitro*, in order to investigate whether the known neurotoxicant metal lead (Pb) impacts the formation of multinucleated microglia. It was determined that isolated rat microglia (Sprague-Dawley), compared to murine (BV2) and human (HMC3) microglia cell-lines, when induced by Phorbol-myristate-acetate (PMA) results in the greatest amount of cell multinucleation and multinuclearity. In this model, it was shown that long-term exposure to Pb inhibited microglia cell proliferation, multinucleation, and multinuclearity in a dose-dependent manner. Pb appears to inhibit multinucleation in microglia by induction of multinuclear regression (karyolysis, pyknosis, or karyorrhexis) resulting in large mononuclear cells. The mechanism by which PMA induces multinucleation may involve p38/MAPK activity and Pb interferes with this activation.

## Introduction

Multinucleated giant cells formed from monoctye/macrophage cells is a feature of numerous pathologies involving inflammation and granulomas (1-3). Microglia play a vital role in many neuro-pathologies, and like other monocyte/macrophage cells have the ability to become multinucleated(4, 5). Multinucleated giant cells in the brain were described as arising from endogenous microglia, as early as 1986, in patients with AIDS related encephalomyelopathy (6). The presence of multinucleated microglia increases with aging, some pathologies, and are inducible by exposure to LPS, beta-amyloid, α- synuclein, TNF-α, interferon-L, and PMA (1, 4). Multinucleated microglia are associated with HIV related dementia and encephalopathy, as well as, tuberculosis and sarcoidosis in the central nervous system (7-9). It has been debated that they form from the fusion of several monocyte/macrophage cells (7, 9), like osteoclasts, but recently it has been demonstrated that multinucleation in microglia occurs through a failure in abscission during cytokinesis (4). Unlike osteoclasts, multinucleation in microglia is reversible(4), and their purpose is unclear. However, due to their larger size, it is speculated that they may be specialized for increased phagocytosis or the ability to engulf other cells (4).

There lacks an abundance of research on multinucleated microglia and previous studies on multinucleation in microglia has been associated with the induction of their formation by infection or inflammatory factors. To date, little investigation into the function of multinucleated microglia has been conducted and there has been no investigation into the role of lead (Pb) on the formation of multinucleated microglia.

Pb is a pernicious metal and environmental contaminant known for its toxic effect on neurons. Recently, our lab has demonstrated a novel role of Pb as a differentiation agent capable of inducing dendritic cell formation and inhibiting osteoclastogenesis, a form of multinucleation, in monocyte/macrophage cells. The formation of dendritic cells came at the cost of osteoclast formation, and since the phenomenon of dendritic cell formation was reproduced in rat microglia, we hypothesized that multinucleation may also be inhibited in microglia by long-term exposure to Pb.

We devised to investigate the role of multinucleation on microglia function and determine whether Pb interferes with this process.

To carry out this investigation we induced multinucleation in several microglia cell types, and monocyte/macrophage cell-line (RAW 264.7) using phorbol myristate acetate (PMA), as was modeled by Hornik et al., 2014. (4). PMA has also been described as stimulating giant multinucleated cell formation in macrophages, in a pathway involving superoxide, p38/MAPK and PKC activation (5, 10). Therefore, in this study, we also examine the effect of Pb on p38/MAPK and PKC pathways, to determine a mechanism of action for Pb’s effect on multinucleation in microglia.

### Material & Methods

Spraque-Dawley rats were purchased from Taconic Biosciences and cell-lines RAW264.7 (ATTC^®^ TIB-71) and BV2 (EOC-2, ATTC^®^ CRL-2469^TM^) were purchased from ATCC. HMC3 cells were obtained from the Reznik Lab at St. John’s University, Queens, New York. All experiments were replicated 3 or more times for optimization and to establish reproducibility. Figures represent graphical representations of reproducible findings.

### Microglia Isolation

Microglia were isolated from the brain of 14-day old Sprague-Dawley rats. Brains were harvested via decapitation and placed on ice in MEM (Corning^®^ cellgro^®)^, containing 1% antibiotics. Tissue was homogenized using a dissociation media supplemented with DNAse I and aspirated with 5 and 10 ml plastic serological pipettes. Dissociated tissue was collected and digestion was halted with ice cold MEM containing 10% FBS and 1% antibiotics. The process was repeated, minus the addition of DNAse I, until all brain tissue was dissociated. Homogenate was pooled and centrifuged at 2000 rpm for 5 minutes, and supernatant was removed. The pellet was re-suspended in fresh growth media (DMEM Corning^®^ cellgro^®^) + 10% FBS + 1% penicillin/streptomycin) and passed through a 40 µm cell strainer before plating in 225 cm^2^ vented tissue culture flasks. A mixed glial culture resulted from this preparation with microglia loosely attached on a bed of astrocytes. Microglia were in suspension and loosely attached to the astrocyte bed and harvested via gentle rocking and washing with cold PBS for 5 to 10 min. After centrifugation cells were resuspended in fresh growth media and purified by sub-culture every 10 minutes for 30 minutes in humidified air containing 5% CO_2_ at 37°C. Microglia were stained with Ib1a antibody to verify identity and determine culture purity.

### Multinucleation Model

To ascertain and validate a suitable model to study multinucleation in microglia, three microglia cell types, and one non-CNS monocyte/macrophage cell-line were compared. Primary rat microglia isolated from Spraque-Dawley rats, murine BV2 microglia cell-line, human HMC3 microglia cell-line, and murine RAW 274.7 monocyte/macrophage cells were stimulated with PMA to induce multinucleation. Cells were incubated with 10, 100, and 1000 ng/ml PMA in MEM supplemented with 10% FBS and 1% penicillin/streptomycin, for 24, 48, and 72 h to determine the optimal concentration and exposure time for multinucleated microglia formation.

### PMA and Pb Co-Exposure Effect on Proliferation

Microglia were plated at a density of 1.5 x 10^5^ cells/well in MEM completed with 10% FBS and 1% penicillin/streptomycin. Cells were incubated overnight in humidified air and 5% CO_2_ at 37°C. Growth media was replaced with completed MEM containing 1000ng/ml PMA only or PMA and Pb (0.01, 0.1, 1.0, 5.0, 10 µM). Control did not receive PMA or Pb, but had completed growth media changed. Cells were further incubated for 24 h before being harvested for counting. Cells were counted on a cell counter (Luna^tm^ Dual Fluourescence) using acridine orange and propidium iodide. Proliferation was determined by subtracting the number of live cells after 24 h from the number of live cells initially seeded. Results from three independent experiments were graphed in comparison to the PMA only treatment.

### PMA Induced Multinucleation and Pb Exposure

Microglia were plated at a density of 1.5 x 10^5^ cells per well in MEM supplemented with 10% FBS and 1% penicillin/streptomycin, and incubated overnight in humidified air and 5% CO_2_ at 37°C. Growth media was withdrawn and replaced with MEM supplemented with 10% FBS, 1% penicillin/streptomycin, 1000 ng/ml PMA, and Pb ( 0.01, 0.1, 1.0, 5.0, and 10 µM). Positive and negative control groups did not receive Pb. Cells were returned to incubation and observed for 72 h for the formation of multinucleated cells.

### Fixing and Nuclei Staining

At 72 h media was withdrawn and cells were washed twice with PBS. Cells were then fixed in well with 10% neutral buffered formalin for 10 min, washed once with PBS, and dehydrated with 50% ethanol. Cells were then stained with 50% Gills hematoxylin for 3 min to visualize the cell nuclei. After removal of hematoxylin, cells were washed with tap water (2X), 0.3% ammonia water (1X), tap water (1X), and rinsed with 50% ethanol. Finally, wells were covered with glycerin before visualizing via light microscopy (Primovert Inverted Microscope, Zeiss).

### Scoring Multinucleation

Each well was divided into 4 quadrants. To count the number of multinucleated cells, the lens was centered in each quadrant and large cells containing more than 2 nuclei were counted as multinucleated. Large cells with disrupted nuclei (indicating nuclear regression) were also counted as multinucleated microglia. The total of all four quadrants were tallied for each experiment (3 independent experiments were carried out for each concentration of Pb, and the negative and positive controls). Wells were blind scored by two investigators.

### Scoring Multinuclearity

The multinuclearity, or number of nuclei per multinucleated cell was also recorded. Each well was divided into 4 quadrants and large cells containing greater than 2 nuclei were considered multinucleated microglia. The number of nuclei per each multinucleated cell counted was recorded. Cells undergoing regression (karyorrhexis, karyolysis, or pyknosis), were denoted as R.

### Nuclear Regression Analysis

Regression was determined by the histological presentation of Karyorrhexis, karyolysis, and pyknosis. Cells undergoing nuclear regression lacked nuclear envelopes and had diffuse hematoxylin staining.

### P38/MAPK Activity Assay

Primary rat microglia cells were seeded at 2.5 X10^5^ cells per well in 6-well plates in growth media (DMEM + 10% FBS +1% penicillin/streptomycin) and incubated in humidified air containing 5% CO_2_ at 37°C overnight. For determination of Pb’s effect on p38/MAPK activity, media was aspirated and replaced with growth media containing Pb (0,0.01, 0.1, 1.0, 5.0 and 10 µM) and cells were returned to incubation for 24 h. Cells were removed from incubation and placed on ice while lysed in media, cell lysis was then used to in p38 activity kit (p38 MAPK (Total) Multispecies InstantOne™ ELISA Kit, Thermofisher Scientific, NY). Colorimetric response was read at an absorbance of 450nM in a microplate reader. For determination of Pb’s effect on p38/MAPK activity in PMA-treated microglia, cells were seeded and incubated overnight as previously described. Growth media was withdrawn, and cells were co-exposed to PMA (1000ng/ml) and varying concentrations of Pb (0, 0.01, 0.1, 1.0, 5.0, and 10 µM) for 18 h and p38/MAPK activity was assessed as previously described.

### PKC Activity Assay

Primary rat microglia cells were seeded at 2.5 X10^5^ cells per well in 6-well plates in growth media (DMEM + 10% FBS +1% penicillin/streptomycin) and incubated in humidified air containing 5% CO_2_ at 37°C overnight. Cell lysate was used in PKC activity kit (PKC kinase activity kit, Enzo Life Sciences, NY). Results of PKC assay were normalized with respective protein concentrations as determined by BCA protein assay.

### Statistical Analyses

All bar graphs are representative of three independent experiments. Data was analyzed using GraphPad Prism 7.04. Unpaired t-test and one-way ANOVA with or with-out multiple comparisons were performed to determine significant differences, with P ≤ 0.05 regarded as significant, and error bars represent the standard error of the mean.

## Results

### PMA Induction of Multinucleation in Microglia

24 h exposure to PMA did not produce any multinucleated cells at any of the tested concentrations or in any of the cell types. However, multinucleated cells appeared at 48 h in all cell types exposed to 10, 100 and 1000 ng/ml PMA. Multinucleation was greatest in treatment groups with concentrations of PMA greater than 10 ng/mL. However, multinucleation did not increase with exposure to concentrations of PMA beyond 100 ng/ml, and the number of multinucleated cells was comparable between 100 and 1000 ng/ml PMA. Additionally, the number of multinucleated cells did not increase with exposure times greater than 48 h. However, 72 h exposure did increase the multinuclearity (number of nuclei per multinucleated cell) in all cell types, except HMC3 human microglia which had optimal multinuclearity between 24 and 48 h. (Figure 1)

**Figure I.**
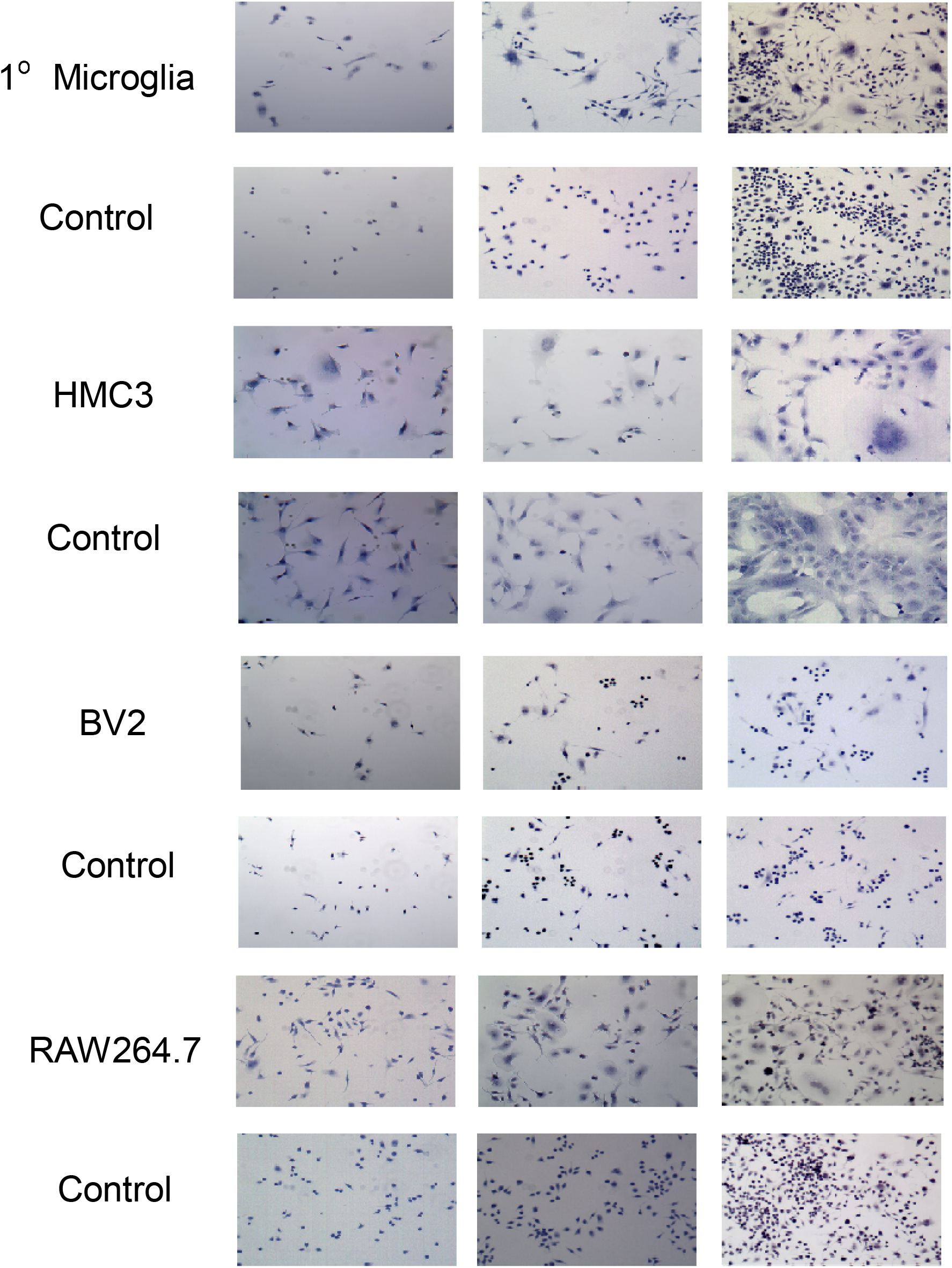
24h, 48h, 72h PMA (100 ng/ml) exposure in 3 microglia cell types and RAW 264.7 monocyte/macrophage cells. Cell population increased over time in both control and PMA exposure groups for all cell types. Large and multinucleated cells appeared by 48 h and persisted or increased up to 72 h. Images acquired at 20x magnification under light microscopy (Primovert Inverted Microscope, Zeiss)

Interestingly, cell size increased before the appearance of multinucleation in all cell types. However, though cells grew larger, in BV2 murine cell-line, microglia resisted multinucleation, and the visibility of distinct nuclei was hampered in HMC3 cells by the lack of both clear nuclear envelops and centrally localized nuclei. Also, HMC3 cells formed sheet-like connections with each other by 72 h, making it difficult to distinguish one cell from another. In contrast, primary rat microglia had clear centrally localized nuclei within visible nuclear envelopes, and produced plentiful large multinucleated cells that persisted for up to 72 h. (figure 1).

It is important to note that a small percentage of microglia are naturally multinucleated, and regression of multinucleation is a natural phenomenon, suggesting that multinucleated microglia are short-lived and develop in response to specific insult. We found that regression occurred in PMA induced multinucleation, increased after 72 h, and was most apparent by 96 h, in primary rat microglia.

### Phagocytic Ability of Multinucleated Microglia

To elucidate the significance of multinucleation, the phagocytic ability of multinucleated microglia was compared to that of mononucleated microglia. Both naive mononucleated and naturally occurring multinucleated microglia were compared to PMA stimulated microglia in their ability to phagocytize small (1.75 µm) and large (6.0 µm) latex beads (figure 2 and 3). Latex beads were chosen for their non-immunogenic characteristics compared to conventional particles used in phagocytosis assays, like zymosan which can activate p38/MAPK pathways (11).

**Figure II.**
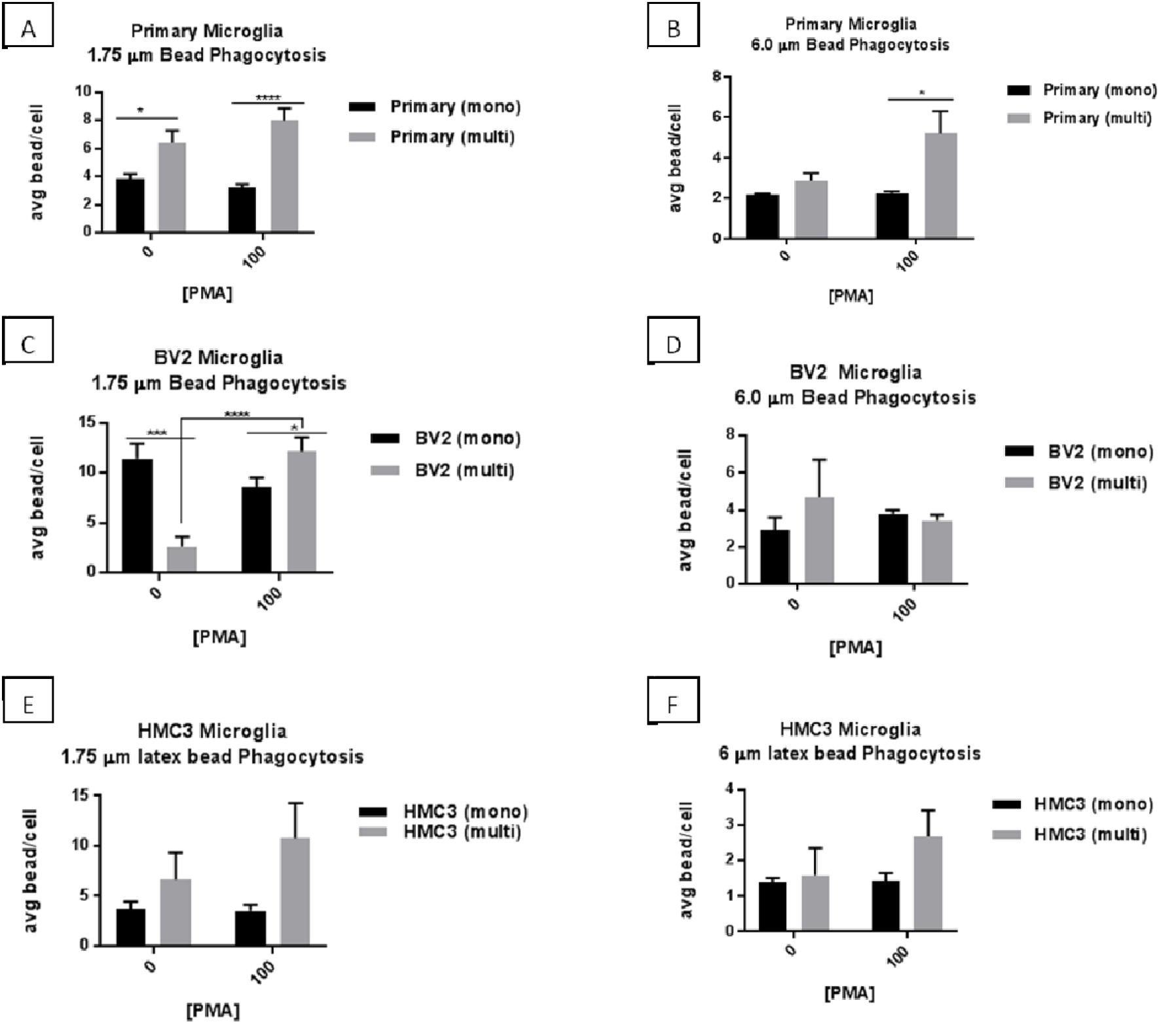
Comparison of Phagocytic Ability in Multinucleated versus Mononucleated Microglia in 3 Microglia Cell Types. PMA induced and naturally occurring multinucleated primary rat microglia phagocytosed a statistically greater number of 1.75 µm beads per cell than their mononucleated counterparts (A). PMA induced multinucleated primary rat microglia had the ability to phagocytose larger beads (6 µm) than PMA mononuclear and untreated mononuclear and multinucleated cells. (B) PMA induce multinucleated BV2 murine microglia had a statistically greater phagocytic ability for 1.75 µm beads than naturally occurring multinucleated BV2 and PMA mononucleated BV2, and mononucleated untreated cells were statistically more phagocytic than naturally occurring multinucleated BV2 (C). There was no difference between the ability of PMA treated BV2 (multinucleated and mononucleated) to phagocytose larger beads (6 µm) than their untreated counterparts (D). HMC3 human microglia did not demonstrate any difference in phagocytic ability between mononucleated and multinucleated cells for 1.75 (E) or 6 µm (F) beads. Graphs represent the average bead/cell score for 3 independent experiments (statistical significance determined by unpaired *t* test. *P ≤0.05 ***P < 0.0005, ****P < 0.0001).

**Figure III.**
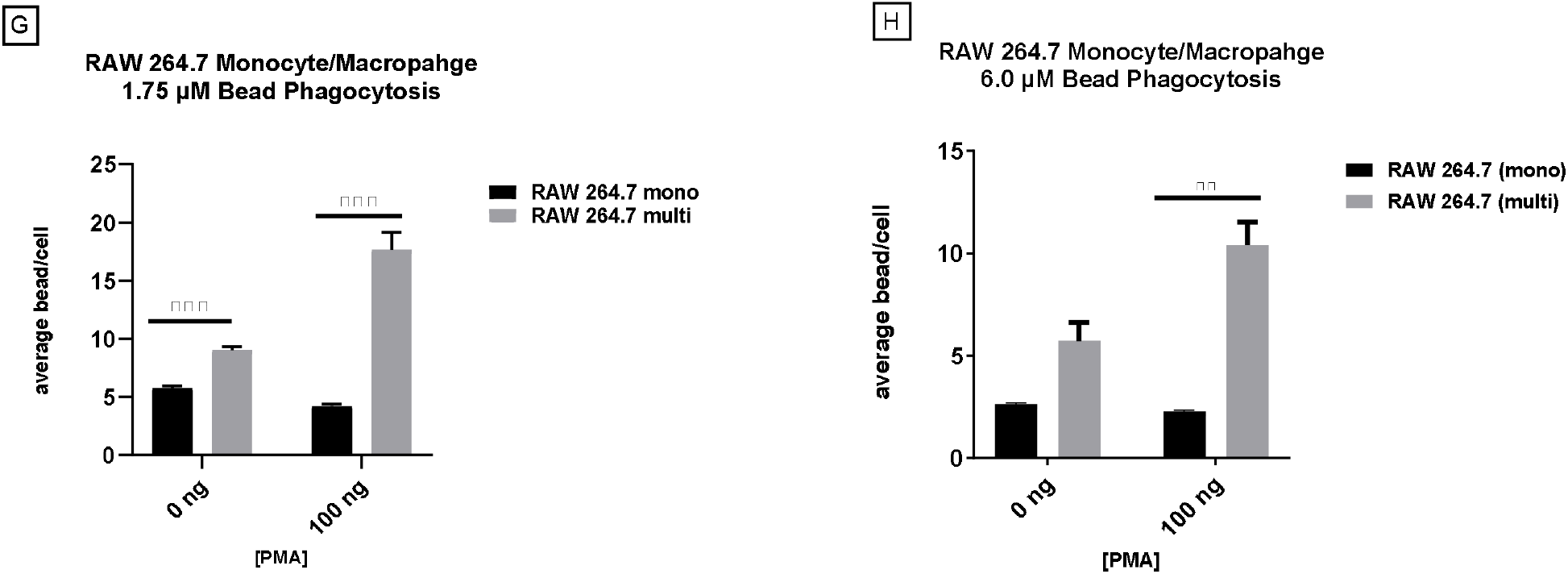
Phagocytic ability of monocyte/macrophage cells (RAW264.7) for small (1.75 um) and large (6.0 µm) particles (latex beads). Both control (naturally occurring) and PMA induced Multinucleated cells had greater phagocytic ability for small particles, compared to mononuclear cells in their respective wells (G). However, for large particles, only PMA induced multinucleated cell displayed a statistically significant increased phagocytic ability, compared to their mononuclear counterparts (H). (statistical significance determined by unpaired *t* test. **P ≤0.005 ***P < 0.0005).

Phagocytosis of small and large latex beads differed between naïve and PMA treated mononucleated and multinucleated primary rat microglia. In naïve (no PMA treatment) primary rat microglia mononucleated cells and naturally occurring multinucleated cells, multinucleated cells phagocytized significantly more small beads than their mononucleated counterparts. Interestingly, there was no statistically significant difference in ability of naïve primary rat multinucleated cells to phagocytose large beads, compared to mononucleated cells. However, phagocytosis of large beads differed between PMA treated mononucleated and multinucleated primary rat microglia. Primary rat microglia that remained mononucleated after being exposed to PMA phagocytized fewer small and large beads, compared to PMA induced multinucleated cells, and these finding were consistent in non-CNS monocyte/macrophage (RAW 264.7 cells) (figure 4). Indicating that PMA induced multinucleation enhances phagocytic ability and points at p38/MAPK and PKC activation, which is involved with the phagocytosis signaling pathway(11), as the mechanism of action. Results differed for the other two microglia cell types (BV2 and HMC3). Strikingly, mononucleated naïve BV2 phagocytized significantly greater small beads than their multinucleated counterparts, but in PMA treated BV2, multinucleated cells took up significantly more small beads than BV2 that remained mononucleated. There were no significant differences amongst any of the comparisons in BV2 for phagocytic ability against large beads (naïve vs PMA, small vs large beads, and mono-vs multinucleated). HMC3 also failed to demonstrated any differences in phagocytic ability between mononucleated and multinucleated cells, whether naïve or PMA treated, for small or large beads.

**Figure IV.**
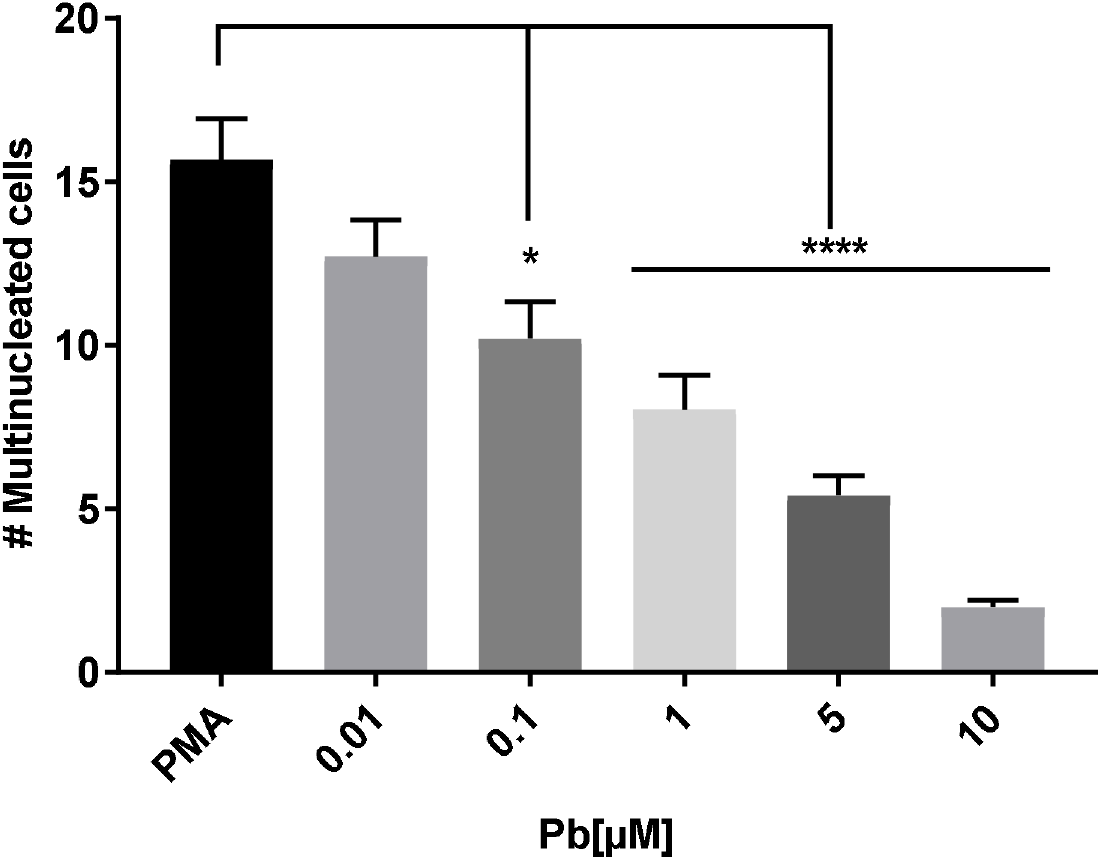
Pb inhibited PMA-induced multinucleated microglia formation. Four quadrants from three independent experiments were scored by two researchers via light microscopy for multinucleated microglia. Large cells containing greater than 2 nuclei and clearly divided nuclear envelopes were categorized as multinucleated microglia. Blind scoring was conducted to avoid bias. Pb dose dependently inhibited PMA induced multinucleation in isolated rat microglia. Graph represents the mean of three independent experiments (statistical significance determined by unpaired *t* test. **P< 0.005, ****P< 0.0001).

### Effect of Pb Exposure on PMA Induced Multinucleation

Long-term co-exposure to Pb and PMA (1000 ng/ml) in primary rat microglia revealed a Pb-related dose-dependent decrease in proliferation (figure 7A), multinucleation (figure 7B), and multinuclearity (figure 6a-f).

### Effect of Pb on Microglia Multinucleation

Long-term Pb exposure dose-dependently inhibited PMA induced multinucleation in primary rat microglia. The number of multinucleated cells (large non-regressing microglia with greater than 2 nuclei) decreased as the concentration of Pb increased, compared to PMA control. Significant inhibition occurred with exposure to 0.1, 1.0, 5.0, and 10 µM Pb for 72 h (figure 4). The decrease seen in the number of multinucleated cells correlated with an overall decrease in the total number of cells.

### Effect of Pb on Microglia Proliferation

Pb decreased proliferation of PMA stimulated microglia after 24 h co-exposure. Pb decreased proliferation in a dose-dependent manner, with proliferation declining in line with increasing concentrations of Pb (figure 7A). Decreased proliferation occurred after 24 h and preceded multinucleation which occurred after 48 h exposure to PMA. When adjusting for population loss the percentage of multinucleated cells present continued to show a Pb-related decrease in multinucleation (figure 7b).

### Effect of Pb on Microglia Multinuclearity

In addition to loss of multinucleation, long-term Pb exposure affected the quality of multinucleated cells formed. Pb exposure resulted in decreased cell multinuclearity (the number of nuclei per multinucleated cell) and increased regression of multinucleation (figure 5a-f, 6a-f, 8). The average number of nuclei per multinucleated cells decreased with exposure to increasing concentrations of Pb, compared to PMA alone. This response was accompanied by regression of multinucleation, which increased with the concentration of Pb. The percent of regressing multinucleated microglia significantly increased with exposure to 5 and10 µM concentrations of Pb (figure 8).

**Figure V.**
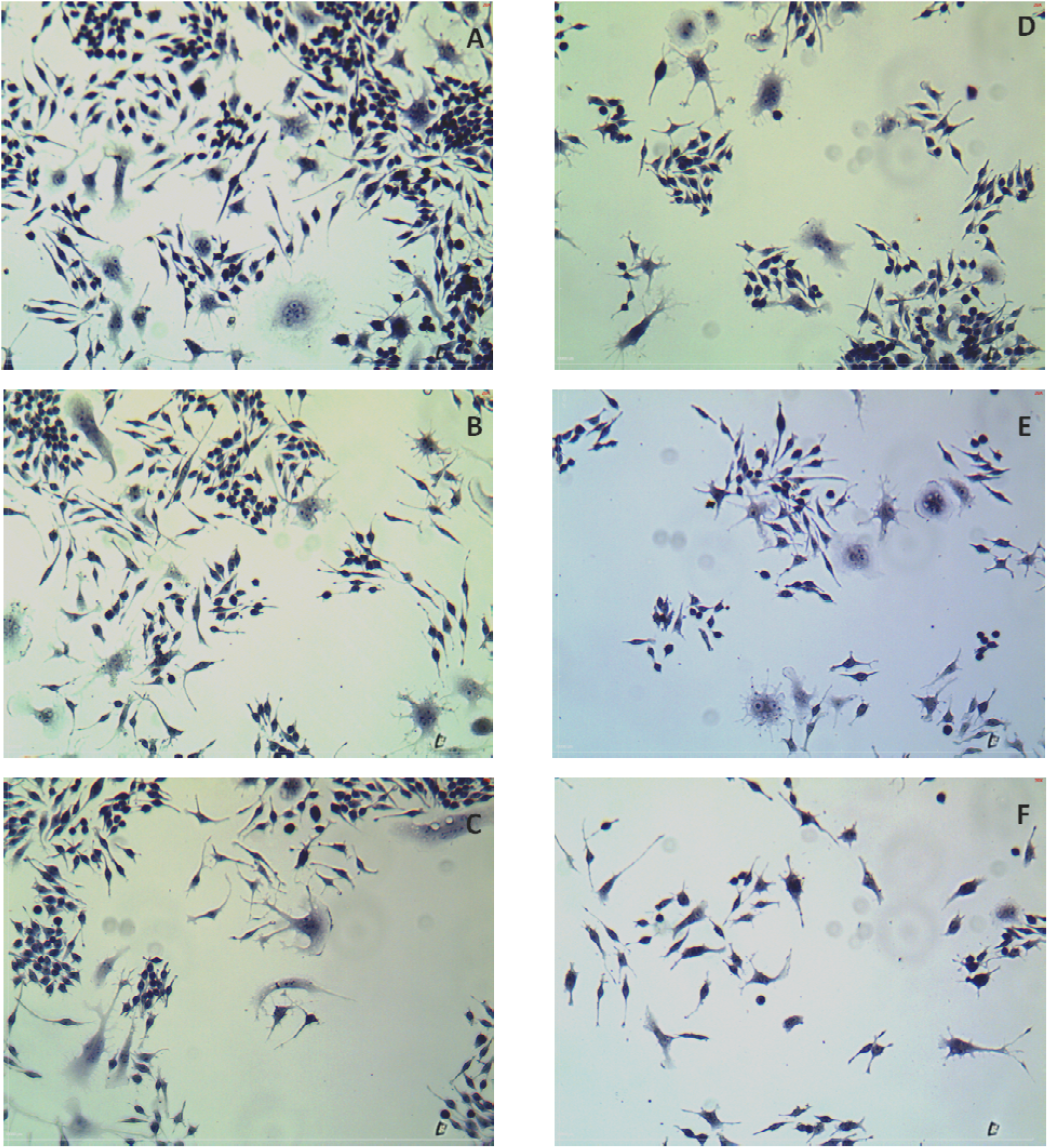
Representative Images of PMA Induced Multinucleation in Pb Exposed Primary Rat Microglia. Large multinucleated cells with distinct nuclei were formed in rat microglia with 72 h exposure to PMA **(A)**. Co-exposure to 0.01 µM Pb had no apparent effect on PMA induced multinucleated cells, and was comparable to PMA positive control multinucleation and multinuclearity **(B)**. Concentrations of Pb 0.1 **(C)**, 1.0 **(D)**, 5.0 **(E)**, and 10 µM Pb **(F)**, induced **karyolysis, karyorrhexis, or pyknosis** in PMA multinucleated cells (disappearance of nuclear envelope and nuclear fragmentation or increased nuclear condensation with cell shrinkage).

**Figure VI.**
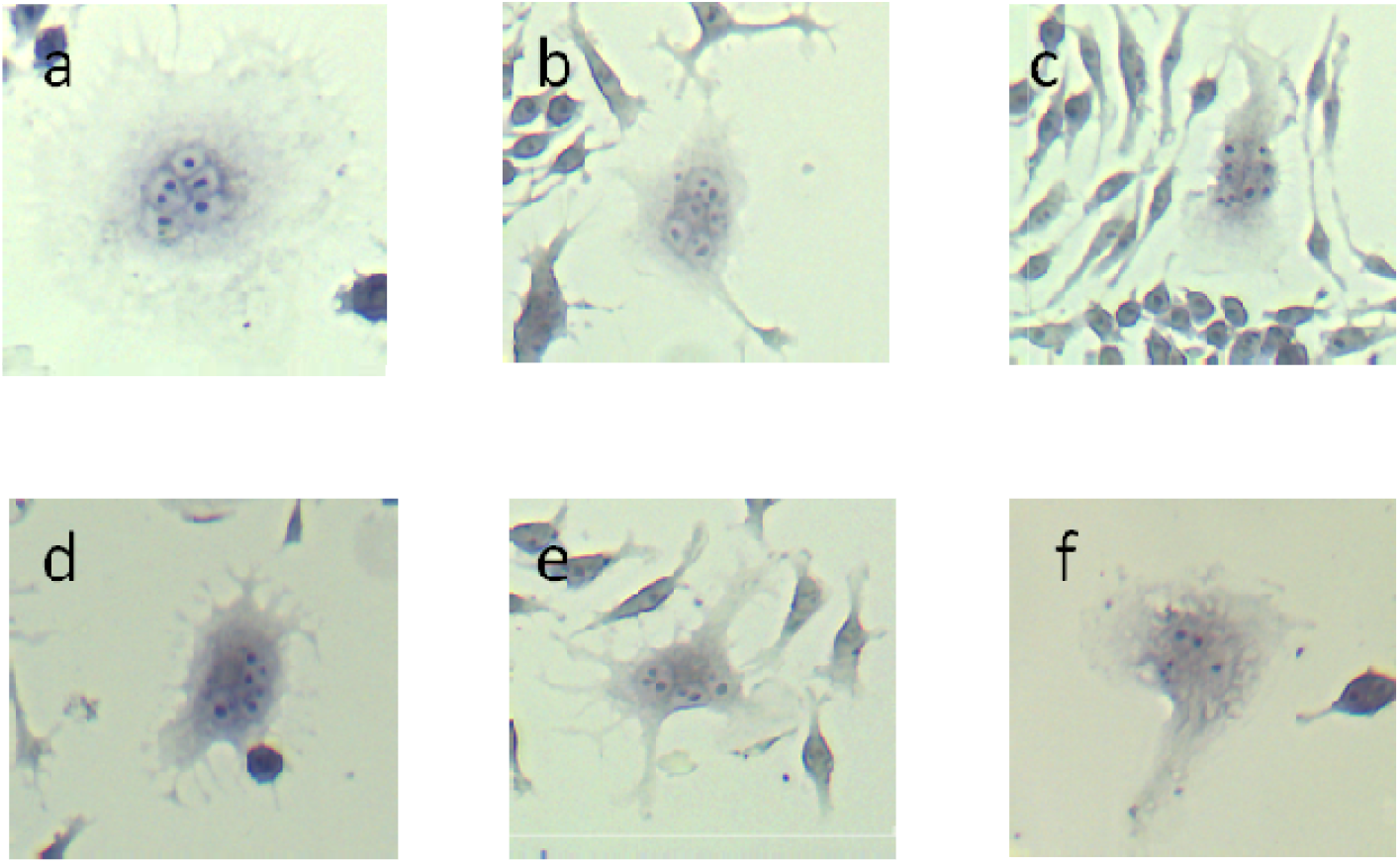
Representative images of multinucleated microglia formation in PMA- treated cells with and without exposure to Pb. **(a)** PMA (1000ng/ml) only, **(b)** 0.01 µM Pb, **(c)** 0.1 µM Pb, **(d)** 1.0 µM Pb, **(e)** 5.0 µM Pb, **(f)** 10 µM Pb. Cells were fixed in 10% neutral buffered formalin and stained with 50% Gill’s hematoxylin. Images were acquired via light microscopy under 20X magnification on a Zeiss inverted microscope with Moticam 3+ microscope camera and Motic Images Plus 3.0 software.

**Figure VII.**
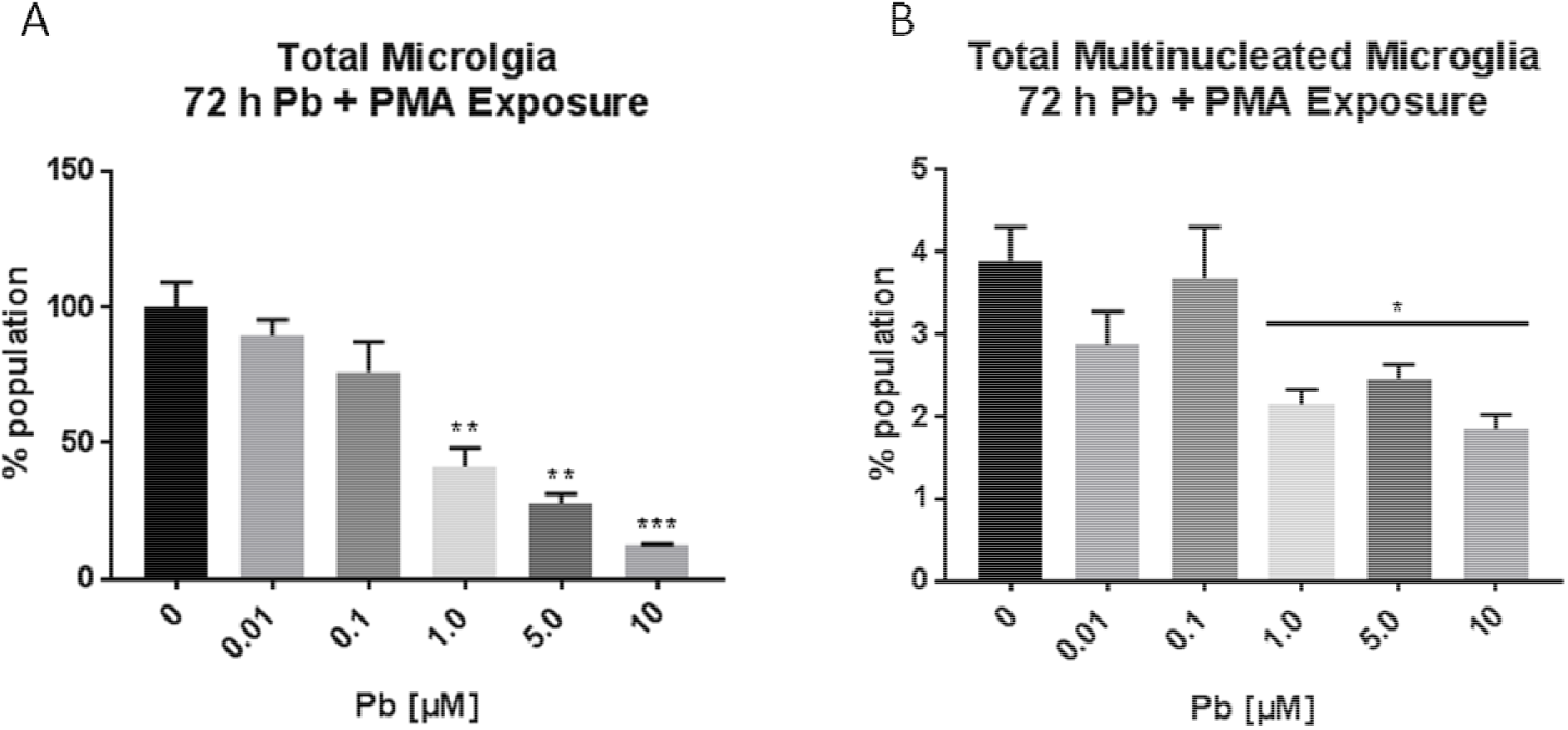
Long-term Pb Exposure Resulted in Decreased proliferation and multinucleation in PMA Exposed Microglia. 72 h Pb exposure dose-dependently decreased the total number of cells. Pb significantly decreased the total microglia population at concentrations ranging from 1.0 to 10 µM compared to control **(A)**. Pb also significantly decreased the total multinucleated microglia population at 1.0, 5.0, and 10 µM concentrations **(B)**. Images were taken using Moticam 3+ camera under 20X magnification and Image Plus 3.0 software. The total number of cells (mononucleated + multinucleated + regressing) and the total number of multinucleated (non-regressing and regressing) was counted using ImageJ cell counter. Data was normalized for percent of population before averages were taken. Bars represent the mean of 3 independent experiments (P* ≤ 0.05, P**<0.01, P***<0.001, Statistical significance was assessed using unpaired parametric t-test).

**Figure VIII.**
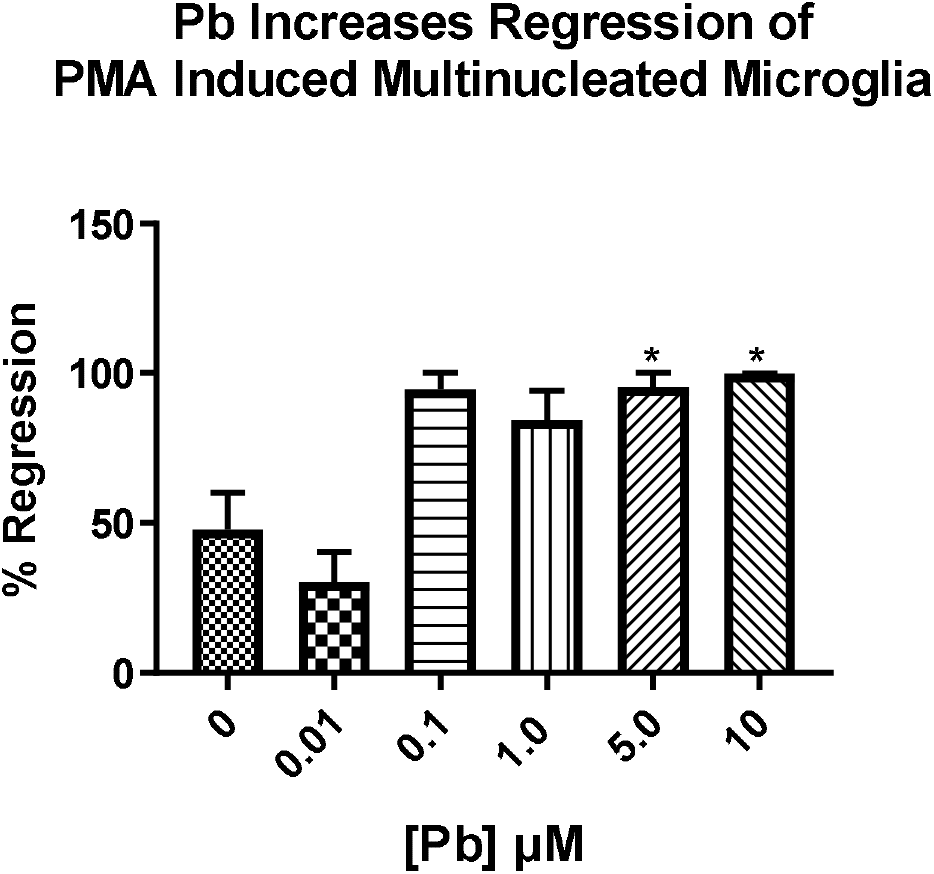
Pb Increased the Percentage of PMA Induced Multinucleated Microglia Undergoing Regression. Co-exposure to PMA and Pb in primary rat microglia resulted in a Pb- associated dose-dependent increase in the number of cells undergoing regression at 72 h. Regression was categorized as any large cell with an obscure nuclear envelope and nuclear fragmentation or pyknosis. Bars represent the mean of three independent experiments (Statistical significance was assessed using unpaired parametric t-test, P*≤0.05).

### Effect of Pb on p38/MAPK Activity in Microglia

There was no significant difference in the activation of p38/MAPK in microglia exposed to concentrations of Pb ranging from 0.01 to 10 µM Pb, with exposure up to 24 h (figure 9). However, in microglia treated with PMA, Pb inhibited PMA-induced p38/MAPK activity. There was a significant decrease in p38/MAPK activity observed with exposure to 0.1, 1.0, and 5.0 µM Pb (figure 10). This indicates that Pb alters cell differentiation through interference with the p38/MAPK pathway.

### Effect of Pb on PKC Activity in Microglia

Pb did not significantly alter PKC activity in rat microglia with 24-hour exposure to concentrations ranging from 0.01 to 5 µM, but a trend indicating an increase in activity as Pb concentration mounts was identified by statistical analysis. As such, a difference in PKC activity was seen with 24 h exposure to 10 µM Pb in which activity was significantly elevated (figure 11).

**Figure IX.**
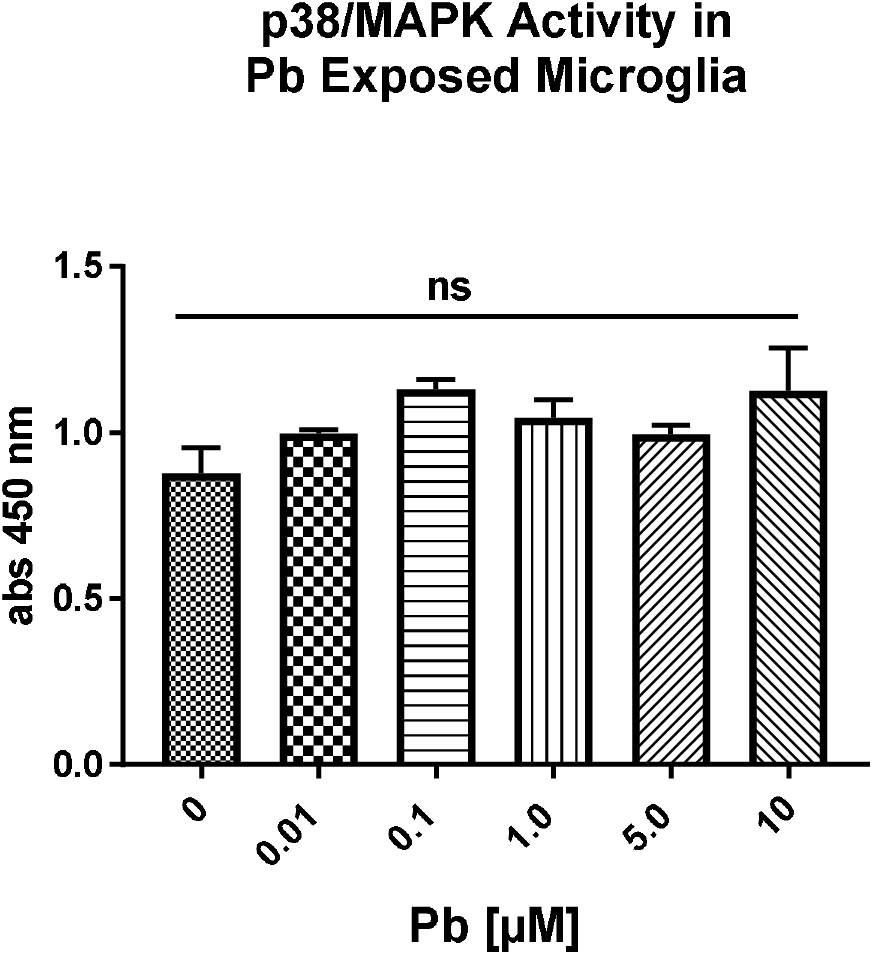
Effect of Pb on p38/MAPK Activity in Rat Microglia. Exposure to Pb (24 h) did not alter p38/MAPK activity in rat primary microglia. There was no significant difference in the activation of p38/MAPK in rat primary microglia exposed to concentrations of Pb ranging from 0.01 to 10 µM compared to control.

**Figure X.**
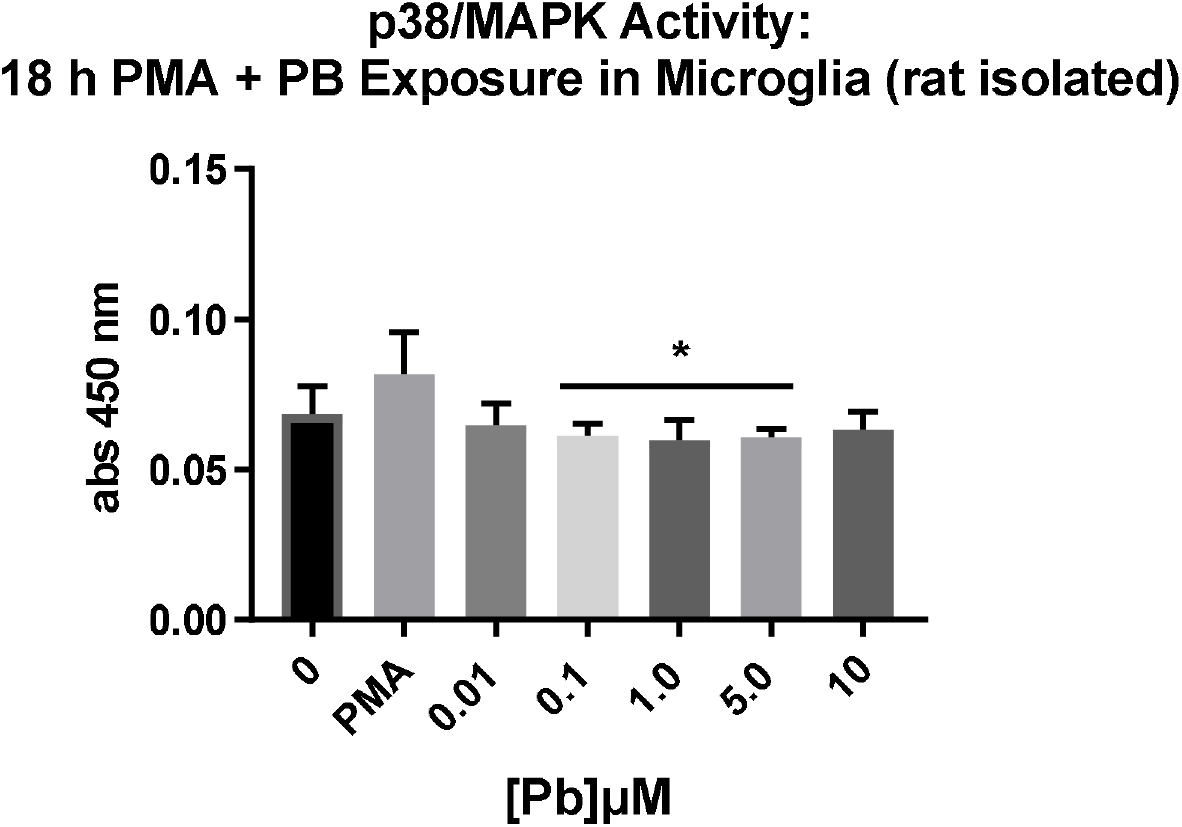
Effect of Pb on p38/MAPK Activity in PMA treated Rat Microglia. Pb exposure (18h) inhibited PMA induced p38/MAPK activity in rat microglia. In primary rat microglia treated with PMA and concentrations of Pb ranging from 0.1 to 10µM, p38/MAPK activity was significantly inhibited in comparison to PMA only treatment. Activity in Pb exposed microglia was comparable to that of cells not treated with PMA or Pb (control). (3 replicates were performed and significant difference was assessed by ANOVA with multiple comparisons, P* ≤0.05).

**Figure XI.**
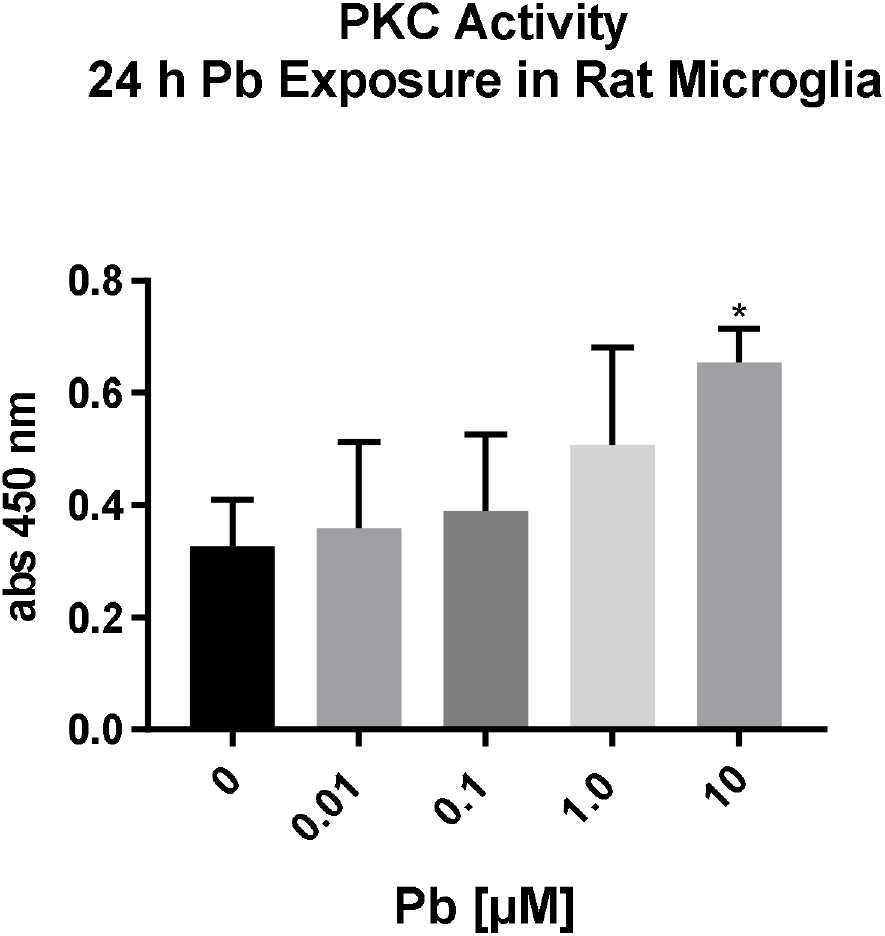
Effect of Pb on PKC Activity in Rat Microglia. PKC activity in primary rat microglia was not significantly altered by 24 h exposure to concentrations of Pb ranging from 0.01 to 1.0 µM. However, there was a significant increase in PKC activity with exposure to 10 µM Pb. (Statistical significance was assessed using unpaired nonparametric t-test, P*≤ 0.05

## Conclusion

It is reported that microglia multinucleation increases with age and in response to pathogenic exposure. However, it is not clear what role they serve; whether their appearance is protective and supportive in the defense and maintenance of the brain homeostatic environment or aberrant and consequential to pathogenesis. Although, it is theorized that multinucleated microglia are specialized to have increased phagocytic capacity and the ability to clear whole cells; possibly due to their increased size (4). To date, few investigations have been reported on the study of multinucleation in microglia cells specifically, while their counterparts in other tissues like the lung have been extensively researched. Therefore, we carried out experiments to determine a suitable *in vitro* model for the study of multinucleated microglia. We compared the ability to induce and to assess multinucleation and multinuclearity in Primary rat microglia, BV2 murine microglia cell-line, and HMC human microglia cell-line, stimulated by PMA. Primary rat microglia exposed to PMA, for 72-hours and at a concentration of either 100 or 1000 ng/mL, was found to be a suitable model for studying multinucleation in microglia; having formed numerous distinct multinucleated giant microglia cells with clearly visible nuclei in a localized region of the cell.

In the present study, multinucleated primary rat microglia were found to have greater phagocytic ability compared to their mononuclear counterparts. Multinucleated Primary rat microglia, both naturally occurring and PMA induced, exhibited greater capacity for phagocytosis of small particles (1.75 µM latex beads). However, only PMA induced multinucleated Primary rat microglia demonstrated increased phagocytic ability for large particles (6.0 µM latex beads). This suggests that multinucleation is induced among microglia cells to increase phagocytic capability (greater number and larger size of latex beads). Furthermore, multinucleation also increased this ability in non-CNS monocyte/macrophage cells, indicating the role of multinucleation in conserved amongst monocyte/macrophage cells whether peripheral or specialized.

Moreover, the PMA/ primary rat microglia model was used to examine the effects of long-term Pb exposure on microglia multinucleation *in vitro*. Long-term Pb exposure had a deleterious effect on microglial multinucleation. Pb inhibited proliferation, multinucleation, and multinuclearity of microglia. Pb exposure inhibited multinucleation through increased nuclear regression of multinucleated microglia. Increased regression was evidenced by the histological appearance of karyorrhexis, karyolysis, or pyknosis; though cell size sometimes remained enlarged.

The mechanism by which Pb inhibits multinucleation was hypothesized to involve p38/MAPK activity. Long-term Pb exposure was previously found by our lab to inhibit RANKL stimulated osteoclastogenesis, the multinucleation of monocyte/macrophage cells, in RAW 264.7 cells, through a mechanism involving the inhibition of p38/MAPK and TNF-α. Pb did not alter the normal activation of p38/MAPK in rat microglia, however it did inhibit PMA stimulated p38/MAPK activity. Furthermore, as PKC interacts with p38/MAPK and is involved in the pathway by which multinucleation and inflammation is regulated in monocyte/macrophage cells, PKC activity was also investigated(12-14). However, PKC activity was not altered by any of the tested concentrations of Pb, except for the highest concentration of 10 µM, in which it was found to be increased. The findings of this study confirm that p38/MAPK activation is required for multinucleation of macrophage cells, and Pb interferes with PMA-induced multinucleation in microglia by inhibiting the activation of p38/MAPK. These findings are interesting, as although PKC is not required for the p38/MAPK formation of multinucleated monocyte/macrophage, it does play a role in accentuating phagocytosis (11, 13).

The phagocytic ability of Pb exposed multinucleated microglia, both regressing and non-regressing, was not analyzed in this study. However, the loss of the ability of microglia to become multinucleated may have consequences for the removal of harmful matter in the brain, as it was demonstrated that multinucleated microglia have increased phagocytic potential. On the other hand, inhibition of proliferation may prevent tissue damage due to inflammation associated with the increased presence of activated microglia and other monocyte/macrophage cells. Further studies both *in vitro* and *in vivo* are required to determine whether formation of enlarged multinucleated microglia is adapted to mitigate inflammation in the CNS while enhancing microglia protective role.

## Conflicts of Interest

There exist no conflicts of interest on the part of any of the authors to declare.

## Notes

### Competing Interest Statement

The authors have declared no competing interest.

